# Identifying systematic variation at the single-cell level by leveraging low-resolution population-level data

**DOI:** 10.1101/2022.01.27.478115

**Authors:** Elior Rahmani, Michael I. Jordan, Nir Yosef

## Abstract

A major limitation in single-cell genomics is a lack of ability to conduct cost-effective population-level studies. As a result, much of the current research in single-cell genomics focuses on biological processes that are broadly conserved across individuals, such as cellular organization and tissue development. This limitation prevents us from studying the etiology of experimental or clinical conditions that may be inconsistent across individuals owing to molecular variation and a wide range of effects in the population. In order to address this gap, we developed “kernel of integrated single cells” (Keris), a novel model-based framework to inform the analysis of single-cell gene expression data with population-level effects of a condition of interest. By inferring cell-type-specific moments and their variation across conditions using large tissue-level bulk data representing a population, Keris allows us to generate testable hypotheses at the single-cell level that would otherwise require collecting single-cell data from a large number of donors. Within the Keris framework, we show how the combination of low-resolution, large bulk data with small but high-resolution single-cell data enables the identification of changes in cell-subtype compositions and the characterization of subpopulations of cells that are affected by a condition of interest. Using Keris we estimate linear and non-linear age-associated changes in cell-type expression in large bulk peripheral blood mononuclear cells (PBMC) data. Combining with three independent single-cell PBMC datasets, we demonstrate that Keris can identify changes in cell-subtype composition with age and capture cell-type-specific subpopulations of senescent cells. This demonstrates the promise of enhancing single-cell data with population-level information to study compositional changes and to profile condition-affected subpopulations of cells, and provides a potential resource of targets for future clinical interventions.

## 1 Introduction

Existing single-cell (SC) gene-expression datasets are typically limited in terms of the number of donors (individuals) owing to high cost of SC technologies. As a result, much of the current research in SC genomics focuses on variation *between* cellular compartments. That is, variation between known populations of cells (primarily cells of different types), which is broadly conserved across individuals. Such studies have been very successful in profiling, for example, cellular organization and localization, cell lineages, and tissue development [1–3]. While studying broadly conserved biological processes from a limited number of individuals is possible in principle due to the expected high consistency across individuals (e.g., Fig. 1a,c), advancing our understanding of heterogeneous conditions that demonstrate molecular variation across individuals requires population-level data. Particularly, probing the *within*-compartment (e.g., cell-type level) etiology of conditions that demonstrate a wide range of effects on gene expression in the population, such as the aging immune system, which is affected by genetic and environmental variation [4], is expected to be very challenging or perhaps infeasible given the small dataset sizes (e.g., Fig. 1b,c; Supplementaty Fig. S1). In such cases, realizing the promise of SC expression to characterize populations of condition-affected cells is therefore expected to require population-level data in order to address inconsistent effects across affected cells of different individuals.

**Figure 1:**
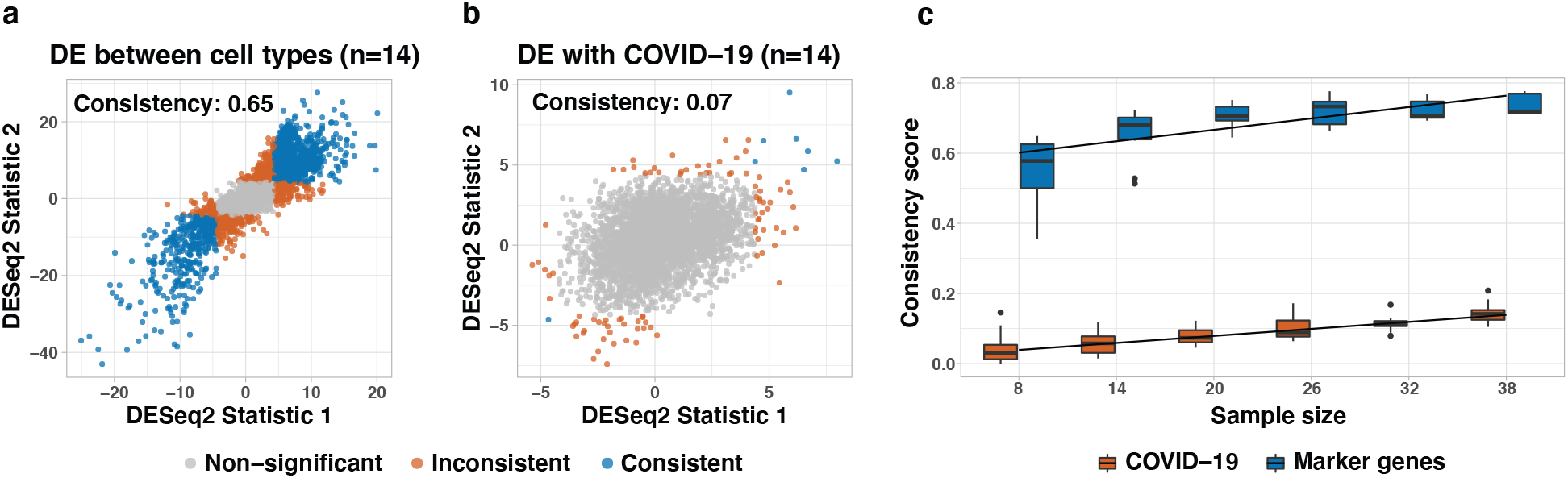
Evaluating the consistency of differentially expressed (DE) genes across two independent PBMC datasets [15, 16]. (a) Test statistics of DE analysis for marker genes *between* main immune cell types in healthy samples, and (b) test statistics of DE analysis between healthy individuals and severe COVID-19 cases, *within* each of the main immune cell types; results from all cell types were pooled together. (c) Repeating the analysis using an increasing number of individuals (sample size); boxplots indicate the performance across 20 randomly sampled subsets of individuals. Consistency was defined as the Jaccard index – the fraction of consistently DE genes (significant in both datasets) out of the total number of DE genes (significant at least once) using DESeq2 [17] on the top 1000 most highly expressed genes, while accounting for age and sex.

In contrast to SC data, tissue-level “bulk” gene expression data effectively aggregates signals over many cells rather than providing a high-resolution cell-specific view of genomics. The relative ease and low cost of generating such data has led to the collection of vast amounts of bulk data from many organisms, tissues, and under different conditions (e.g., bulk profiles from over two million individual samples publicly available on the Gene Expression Omnibus alone [5]).

An understanding that the relative merits of SC and bulk data complement each other has led to the development of methods that leverage SC data as a reference for decomposing heterogeneous bulk profiles in the learning of cell-type compositions [6–12]; this approach can be applied to large datasets for detecting changes in tissue composition that are often demonstrated in disease, such as in cancer [13] and diabetes [14]. Importantly, these methods implicitly rely on the fact that cell-type marker genes are highly consistent across individuals (Fig. 1a,c), which in principle renders SC data a consistent reference for learning cell-type compositions, regardless of the number of individuals in the data. Yet, small SC reference datasets are unlikely to represent well the wide range of effects that different experimental or clinical conditions may have on gene expression within a cell type (Fig. 1b,c). An important question is therefore whether we can change the direction taken by existing methods. That is, can we inform the analysis of SC data with population-level information from large bulk datasets? Such a capability can markedly increase our ability to profile the expression landscape of heterogenous conditions and to characterize populations of cells that are affected by consistent variation in conditions of interest.

### Contributions

We developed “kernel of integrated single cells” (Keris), a novel model-based framework to inform the analysis of SC expression data with population-level variation. We use Keris to infer cell-type-specific moments for the expression of genes and their statistical dependence on experimental or clinical conditions, using tissue-level (bulk) data from large cohorts. This allows us to generate testable hypotheses at the SC level that would otherwise require collecting SC data from a large number of donors. Particularly, we facilitate two applications under the Keris framework, both of which require only typical (i.e., small) SC data given the information extracted by Keris from population-level bulk data. First, the detection of changes in the composition of cell subtypes with a condition of interest, and second, the identification and study of linear and non-linear (gene-gene interactions) statistical effects on expression at the cell level, which, in turn, allow to profile populations of cells that are affected by the condition of interest. We demonstrate our approach in the context of aging and show that Keris enables the identification of changes in cell-subtype composition with age. Further, we use a total of four independent datasets to provide multiple evidence that Keris can identify and enable downstream analysis of unknown subpopulations of senescent cells.

### Related work

In the first part of this work, we propose a model for linear and non-linear effects of a condition on the variation of expression at the single-cell level, and we show that learning the model effectively reduces to learning population-level cell-type-specific co-expression networks (including means and variances) from large tissue-level bulk data. Methods for the construction and detection of co-expression networks [18] and changes in gene-gene covariances (correlations) [19] from bulk data have been previously suggested. However, co-expression based on bulk data primarily captures cell-type composition variation rather than intra-cell-type co-expression signals [20]. The classical decomposition approach with expression data infers only cell-type level means [21], effectively making an unrealistic assumption that all individuals share the same transcriptome at the cell-type level. An understanding that learning cell-type level covariances requires to model the variation of cell-type level expression across individuals has led to another class of methods that aim at explicitly modeling individual- and cell-type-specific variation. However, existing methods do not practically allow the construction of reliable population-level co-expression networks at a cell-type-specific resolution – owing to restrictive assumptions, such as requiring for each individual multiple independent expression measurements [22], or owing to unrealistic assumptions that expression levels from a few individuals represent the population-level distribution of expression [23]. Finally, approaches for estimating individual- and cell-type-specific variation from a single bulk measurement exist [9, 24], however, these do not model co-expression, and as we show later, perform poorly in estimating co-expression.

## 2 Methods

### 2.1 From a single cell to tissue-level variation

#### A proposed model for single-cell expression

Let *Z*_*ijhs*_ be the single-cell expression of individual *i* ∈ {1, …, *n*} in gene *j* ∈ {1, …, *m*}, cell type *h* ∈ {1, …, *k*}, and cell *s* ∈ {1, …, *N*_*ih*_}, where *N*_*ih*_ denotes the number of cells of type *h* coming from a specific tissue under study in individual *i*. We model *Z*_*ijhs*_ in the context of a heterogeneous binary condition *y* (e.g., case/control for a certain disease) and denote the condition of individual *i* by *y*_*i*_ ∈ {0, 1}. We make the following assumptions:

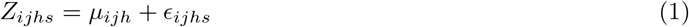

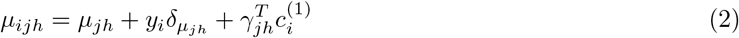

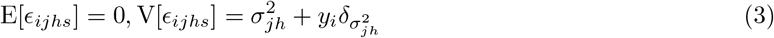

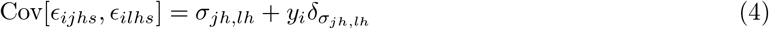

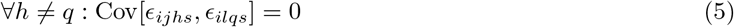

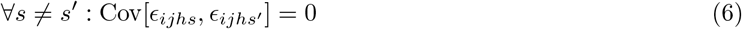

Eq. (1) is comprised of two terms as follows. First, a mean effect *μ*_*ijh*_, described in Eq. (2), which is assumed to have a component specific to gene *j* and cell type *h* (*μ*_*jh*_) and two additional types of possible effects: gene- and cell-type-specific effect 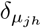 if *y*_*i*_ = 1 and systematic changes in mean, according to *p*_1_ (known) individual-specific factors 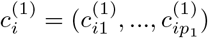 and their corresponding fixed effect sizes *γ*_*jh*_ = (*γ*_1*jh*_, …, *γ*_*p jh*_). Second, a component of variation *ϵ*_*ijhs*_, described in Eq. (3), which is assumed to consist of a systematic component of variation 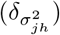 if *y*_*i*_ = 1 and a gene- and cell-type-specific intrinsic variation (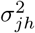; e.g., owing to unmodeled factors, such as the individual’s genetic background and environmental exposures).

Following Eq. (4)-(6), the component of variation *ϵ*_*ijhs*_ may further covary with *ϵ*_*ilhs*_ (i.e., different genes in the same cell). This covariance is assumed to have a gene- and cell-type-specific background covariance (*σ*_*jh,lh*_) with an additional possible effect on the covariance 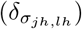 if *y*_*i*_ = 1. Lastly, Eq. (4)-(6) also introduce the assumption of statistical independence between different cell types and between different cells. Of note, despite the assumption of independence between cells, the assumption of cell-type-specific mean profiles in Eq. (2), as well as the covariance structure defined in Eq. (4), impose the expected similarity within populations of cells of the same type by the similarity of their joint distribution across genes.

#### Relating the single-cell model to tissue-level bulk expression

We wish to learn the magnitude of 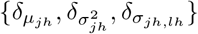 with respect to {*μ*_*jh*_, *σ*_*jh*_, *σ*_*jh,lh*_}, which would essentially allow us to identify changes in moments of expression (means, variances, and covariances). As we show next, under our proposed model, these moments are expected to be reflected in tissue-level bulk expression as well. Establishing this link will eventually allow us to develop inference that can be applied to bulk data. We define a characteristic cell-type level expression of individual *i* in gene *j* and cell type *h*, denoted by *Z*_*ijh*_, as a random variable that preserves the moments of the expression in gene *j* across all cells of type *h* in individual *i*, up to a scaling factor, which is expected to be dominated by the data sampling process (e.g., library size):

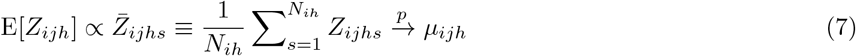

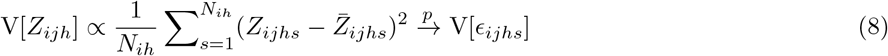

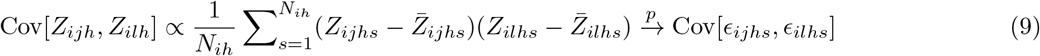

Given the definition of a cell-type level expression, we can now summarize them to get a representation of the tissue-level bulk expression of individual *i* in gene *j*:

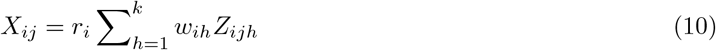

Here, *w*_*i*_ = (*w*_*i*1_, …, *w*_*ik*_) represent the individual-specific cell-type proportions of the *k* cell types composing the tissue under study and *r*_*i*_ is an individual-specific scaling factor. Normalization methods aim at eliminating individual-specific effects of library size [25]. Successfully accounting for these biases would remove the need for including a scaling factor in the model, and we therefore omit it in what follows; we get:

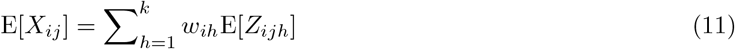

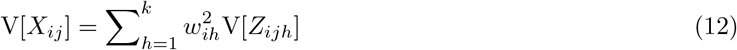

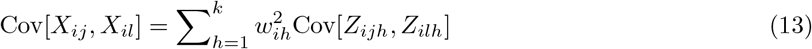

Since the number of cells in a sample is typically very large (neglecting low-frequency cell types), the characteristic cell-type expression levels are expected to reflect the parameters of the model in Eq. (1)-(4) well, and as a result, the bulk levels {*X*_*ij*_} are also expected to reflect them well.

### 2.2 The Keris framework

#### The Keris model for tissue-level bulk expression

We consider the derivation in Eq. (11)-(13) to define our full, distribution-free model for tissue-level bulk expression:

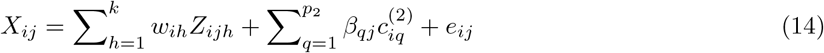

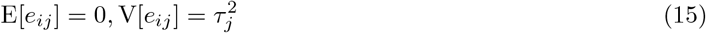

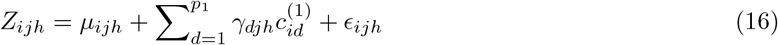

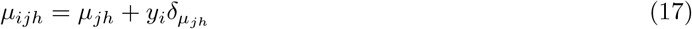

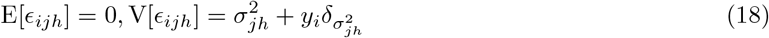

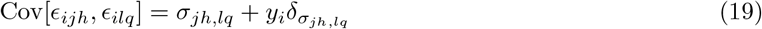

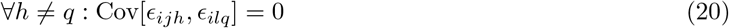

Here, *e*_*ij*_ is an i.i.d. component of variation that reflects measurement noise and 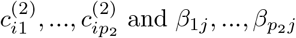 and *β*_1*j*_, …, *β*_*p j*_ are *p*_2_ (known) individual-specific factors and their corresponding fixed effect sizes. The latter model systematic changes in means at the mixture level, which can be the result of the experimental design. Most notably, batch effects and other technical variation may globally affect the mixtures {*X*_*ij*_} regardless of cell-type-level expression; such factors can be estimated by methods for removal of unwanted variation [26, 27]. The covariates 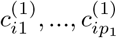,on the other hand, represent factors thay may have systematic effects on mean expression at the cell-type level, such as demographics or clinical status. We work under the assumption that all covariates and the cell-type proportions {*w*_*ih*_} are known. In practice, cell-type proportions can be estimated computationally using existing methods (e.g., [6, 8–10]).

The derivation of the model based on the single-cell perspective in Eq. (1)-(6) renders Keris as the result of an implicit generative model for single-cell expression, and the parameters in Eq. (1)-(4) can be estimated by fitting the model in Eq. (14)-(20) using tissue-level bulk data. Since we model gene-gene covariances, learning the model from large bulk data effectively results in population-level cell-type-specific co-expression networks and their changes with a condition of interest. In the remainder of this Subsection, we discuss the second part of the Keris framework: how combining single-cell data with such population-level co-expression networks can identify changes in composition of cell subtypes and profile subpopulations of cells that are differentially expressed (DE) with a condition of interest. This will be followed by a description of the model inference and statistical testing for differential moments (DM). An Illustration of the Keris framework is given in Fig. 2.

**Figure 2:**
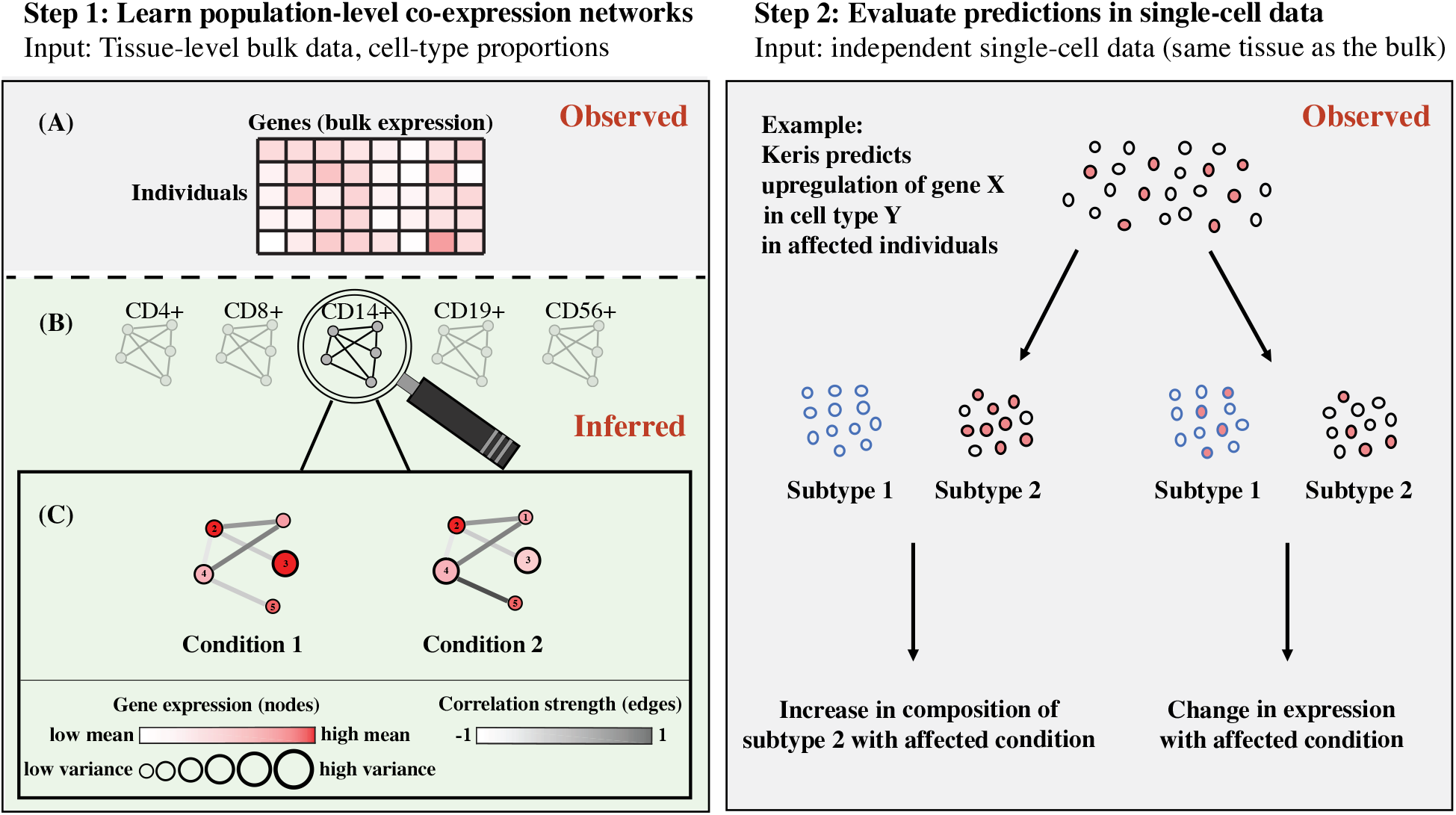
Illustration of the two main conceptual steps in the Keris framework. Step 1 (left): (A) Given bulk gene expression and matching estimates of cell-type proportions, (B) the model infers cell-type-specific coexpression networks, (C) which are then further deconvolved by statistically testing for differential moments (means, variances, and covariances between genes) with a condition of interest (which results in condition-specific networks). Step 2 (right): given single-cell data, we can test whether a differentially expressed gene (or a differentially correlated pair of interacting genes) reported in Step 1 is a marker gene of a cell subtype. This allows us to classify the gene (or the gene-gene interaction) as either indicating a change in cell-subtype composition or indicating a change in expression with the condition under test. Genes and gene-gene interactions that indicate changes in expression can then be used in a single-cell downstream analysis.

#### Detecting variation in cell composition

A real DM signal in a particular cell type can reflect either a change in expression within that cell type or a change in the composition of cell subtypes (or both). As an example, consider testing a case/control status for association with some cell type 𝒞 that represents a class of two cell subtypes 𝒞_𝒜_, 𝒞_ℬ_. An increase in the composition ratio between 𝒞_𝒜_ and 𝒞_ℬ_ in cases versus controls will make genes that are overexpressed in 𝒞_𝒜_ compared with 𝒞_ℬ_ (i.e., regardless of case/condition status) to appear as overexpressed in the group of cases when looking at the heterogeneous class 𝒞.

Naturally, capturing cell-type level signals from tissue-level bulk data is limited to the main cell types in the tissue under study. As a result, DM detected by the Keris model can be due to changes in composition of unmodeled cell subtypes. However, we can use SC data to deduce whether an observed DM signal is the result of changes in cell-subtype composition by evaluating whether a putative DM is marker for a specific cell subtype (Fig. 2). For example, a Keris-derived DM result indicating upregulation with a condition in cell type 𝒞 would imply an increase (decrease) in the composition of cell subtype 𝒞_𝒜_ with the condition if the DM is an upregulated (downregulated) marker for subtype 𝒞_𝒜_. Importantly, given that markers of cell types and subtypes are expected to be highly consistent across individuals (e.g., Fig. 1a,c), this step does not require population-level data and can rely on small SC data.

#### Characterizing condition-affected populations of cells

Eq. (1)-(6) model cell-level changes in expression under a condition of interest *y*. In reality, it is likely that only a subset of the cells of an affected individual (*y*_*i*_ = 1) will be affected by the condition. Further, in general, cells from both groups defined by *y* may demonstrate affected cells. For example, consider the case where *y*_*i*_ = 1 indicates an aged individual, compared with a younger background population. In this case, only a subset of the cells of a given individual are excepted to be aged (senescent), with an expected higher fraction of such cells in older individuals.

Let *Y*_*ihs*_ be a Bernoulli random variable representing the cell-specific senescence state and let *y*_*ihs*_ be its realization, where *y*_*ihs*_=1 indicates a senescent cell, consider the assumption:

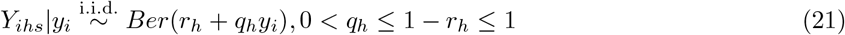

Put in words, a given cell is more likely to be senescent if it is coming from an aged individual. Taking this assumption renders the model in Eq. (1)-(6) unchanged, with the exception that now the parameters 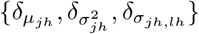 contribute their effects only in cells for which *y*_*ihs*_ = 1. Consequently, our definition of the characteristic cell-type level expression from Eq. (7)-(9) now satisfy:

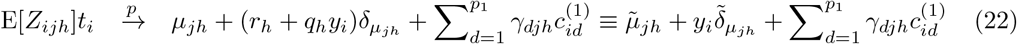

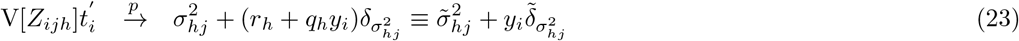

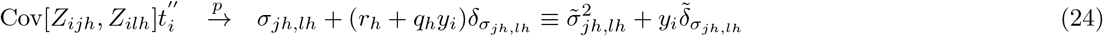

where 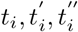 are individual-specific scaling factors.

This perspective shows why non-zero effects 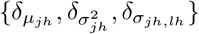 may indicate cellular-level statistical associations with the condition *y*. Moreover, under this view, 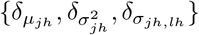 are effects in senescent cells regardless of whether the cells are coming from an aged (*y*_*i*_ = 1) or young (*y*_*i*_ = 0) individual. This implies that a DM identified by Keris can reflect an increased (or decreased) expression in senescent cells compared with non-senescent cells within a given individual - regardless of whether the individual is young or aged.

Put differently, a hypothesis generated by Keris from population-level data can be casted as a hypothesis on a population of cells. As a result, in the case of our example with aging, we can in principle evaluate hypotheses generated by Keris using a population of senescent and non-senescent cells from a single donor (or a few donors) without the need for data from a large number of donors.

### 2.3 Inference and statistical testing

#### Technical background

The Generalized Method of Moments (GMM) allows us to learn parameters of a model based on moment conditions that match population moments with their data-derived sample counterparts [28]. More generally, each moment condition may match a function of multiple parameters of the model with its sample-based estimate (i.e., rather than a moment condition per a single parameter). Estimating the model’s parameters then effectively reduces to solving a system of equations with multiple variables. However, unlike in the Method of Moments, which essentially provides a (unique) solution only in cases where the number of variables equals to the number of equations (assuming full rank), the GMM framework addresses estimation in the case where the number of equations is greater than the number of variables (i.e., an over-determined system of equations). More concretely, let *f* (Θ, *X*_*n*_) = *f*_1_(Θ, *X*_*n*_), …, *f*_*d*_(Θ, *X*_*n*_) be *d* moment conditions defined over the set of parameters Θ of a given model, where *X*_*n*_ = (*x*_1_, …, *x*_*n*_) are observations coming from the model, if the moment conditions satisfy ∀*i* ∈ {1, …, *d*} : E[*f*_*i*_(Θ, *X*_*n*_)] = 0 and certain minimal regularity conditions are met (errors are stationary and ergodic martingale difference sequence) then an asymptotically consistent estimator of Θ for the case *d >* |Θ| is given by [28]:

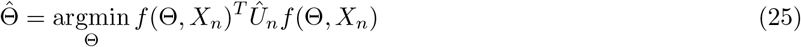

where *Û*_*n*_ is a positive definite (PD) weighting matrix. In the common case of a linear model, Eq. (25) reduces to solving a generalized least-squared problem (GLS). The weights *Û*_*n*_ provide a mechanism to address differences in magnitude of noise in different samples (heteroskedasticity) or other properties of experimental design, such as sampling bias. Most commonly, *Û*_*n*_ assigns the moment conditions with weights so that the estimation is less sensitive to conditions with high variance. For a comprehensive treatment of GMM see, for example, the textbook by Hayashi [29].

#### Optimization setup

Estimation under the original GMM framework relies on large sample properties that arise in the theoretical regime of a constant number of moment conditions and an infinite number of observations that are used for the evaluation of each moment condition. Other formulations of GMM further concern the specific case of multiple equations (i.e., models), which may have common and/or equation-specific parameters. Here, we consider the case where data points of different observations are coming from non-identical distributions that are parameterized by a combination of known observation-specific factors and a set of parameters that are shared across the distributions. In such a scenario, the sampling distribution of each data point has potentially unique population moments, however, the population moments of all the sampling distributions are tied together by a set of common parameters.

More concretely, in the Keris model, the means, variances, and covariances of each data point (or pair of data points) are functions of the parameters of the model and individual-specific factors, including the individual’s cell-type proportions and covariates. This allows us to develop our model optimization by following the concept of GMM estimation under a special regime, wherein each moment condition is calculated from a single observation that represents a single distribution, however, the number of moment conditions (=the number of observations) is infinite, and the parameters defined in the moment conditions are common to all moment conditions. Importantly, while the space of parameters to be estimated is large, as described next, our optimization scheme avoids overfitting by reducing the problem to solving a large number of independent weighted least-squares problems (WLS) problems, such that each problem considers all the samples in the data, yet, only learns a constant number of parameters.

#### Estimating the means in the model

We first develop an estimator for the parameters encoding mean effects in Eq. (14) and (17). We denote 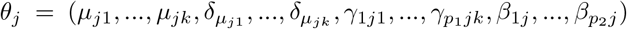 to be the gene-specific mean parameters for a gene *j* ∈ {1, …, *m*}; following our model, this set of parameters can be learned for each gene independently. Let 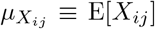 be the individual-specific mean of sample *i* ∈ {1, …, *n*}, we represent *θ*_*j*_ by a vector of *n* moment conditions *f* (*θ*_*j*_) = *f*_1_(*θ*_*j*_), …, *f*_*n*_(*θ*_*j*_) as follow:

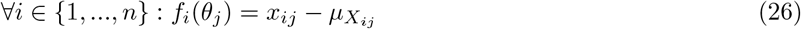

In words, *x*_*ij*_ is used as a single-sample estimate of 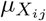 based on the observed value *x*_*ij*_ coming from *X*_*ij*_; clearly, E[*f*_*i*_(*θ*_*j*_)] = 0. Note that *f*_*i*_(*θ*_*j*_) is a function of *θ*_*j*_, *x*_*ij*_, and the other individual-specific factors; we make this compact notation for readability. Denoting *s*_*i*_ as the vector of individual-specific factors 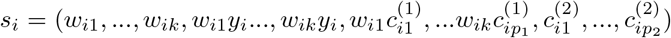 and denoting *S* to be a matrix with *s*_1_, …, *s*_*n*_ stacked as its rows, we define our estimator of *θ*_*j*_ by following Eq. (25) and (26):

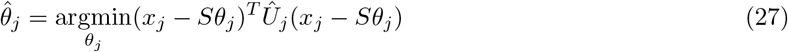

where *x*_*j*_ = (*x*_1*j*_, …, *x*_*nj*_). We can set *Û*_*j*_ ∈ ℝ^*n×n*^ to be the empirical variance of *f* (*θ*_*j*_), which results in a diagonal matrix due to the assumption of independence between individuals:

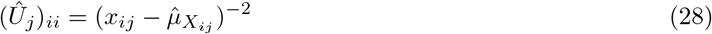

We get a WLS problem, for which the solution and its asymptotic distribution are known:

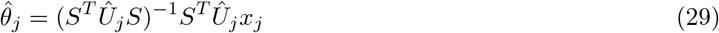

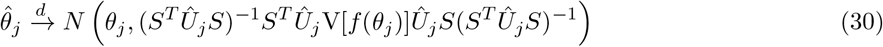

In practice, we can apply a feasible GLS procedure, in which we first set *Û*_*j*_ = *I*_*n*_ and then following a solution of Eq. (27) we can re-estimate Ûbased on Eq. (28) to be used in a subsequent optimization of Eq. (27). While this would result in asymptotic efficiency of the estimator, we make a practical adjustment to the weights in Eq. (28) since each *f*_*i*_(*θ*_*j*_) is based on a single data point. Specifically, in the likely case of a substantial deviation of a given data point from its expected value per the model, following Eq. (28), such a data point would be assigned with a low weight, which would result in a desired down-weighting of this observation in the optimization. However, the reverse case is more of concern: if a given data point demonstrates a high correspondence with the expected value then a subsequent iteration will up-weight it, and for a near-perfect correspondence between a data point and its expected value (which is likely to happen for some data points in large data), its extreme weight can dominate the solution and prevent the optimization from obtaining a good solution across the data. We therefore use the following weighting scheme:

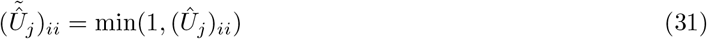

Following Eq. (30) this results in an asymptotically consistent estimator that follows:

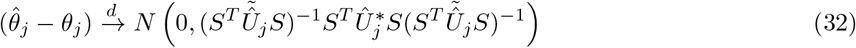

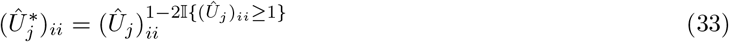

#### Estimating the variances and covariances in the model

We develop estimators for the variances and covariances in the Keris model by following the same concepts we applied for estimating the means. For every pair of genes *j, l* ∈ {1, …, *m*}, we define a vector of parameters as follows:

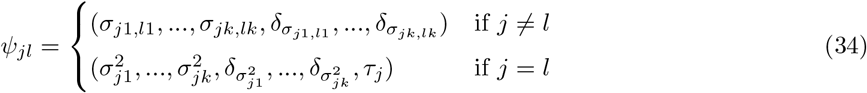

Given 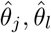, the parameters in *ψ*_*jl*_ can be learned separately from the rest of the parameters in the model. Let Σ_*i*_ ∈ ℝ^*m×m*^ denote the variance of sample *i* ∈ {1, …, *n*} such that (Σ_*i*_)_*jl*_ = Cov[*X*_*ij*_, *X*_*il*_], we define:

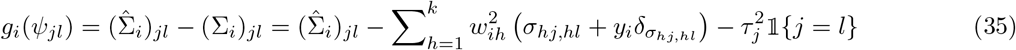

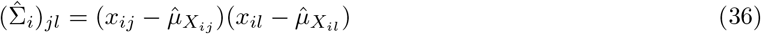

Based on these moment conditions, and similarly to the estimation of the means, we set our estimator:

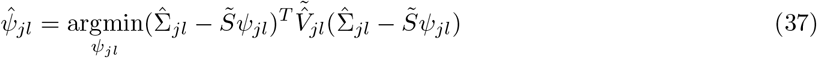

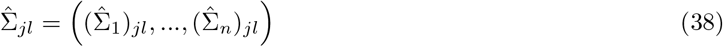

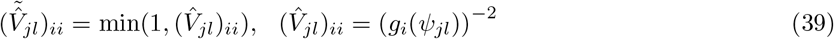

where 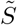 is a matrix with its *i*-th row defining the known factors for the covariance in sample *i*:

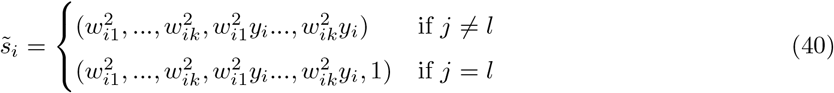

Finally, we construct the estimator and its asymptotic distribution:

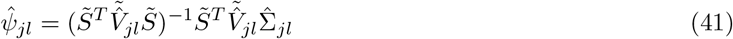

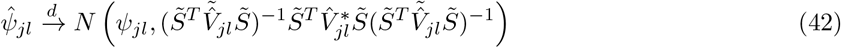

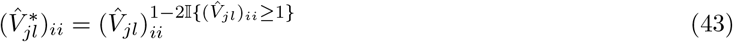

While this estimator is asymptotically consistent, in order to improve estimation in finite data we further consider natural constraints of the problem with the goal of providing more information to direct the inference towards a good solution. Particularly, we constrain the variances to be non-negative, and in the cases where *j* ≠*l*, we consider the following necessary conditions resulting from the definition of covariance:

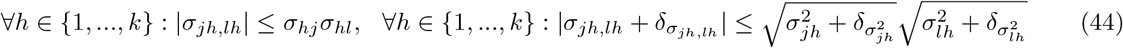

These constraints require estimates of the variances 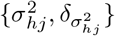. The estimation of the variances does not depend on the covariances in the model (due to the assumption of independence between cell types), and therefore we can learn 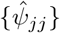 first (i.e., after learning 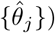) and only then estimate 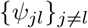.

### 2.4 Statistical testing for differential moments

#### Deriving p-values under asymptotics

The Keris model estimates the statistical effects of a binary condition of interest *y* on the means, variances, and covariances. Given such estimates, a key question of interest is to quantify the statistical evidence for whether each of the effects 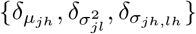 is non zero. Deriving asymptotic t-ratio statistics for the Keris estimators is possible by following Slutsky’s Theorem. The ratios between (I) the estimators that converge in distribution to normal distributions and (II) the estimators of their variances that converge in probability allows for a straightforward statistical testing:

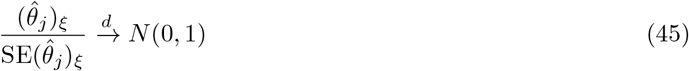

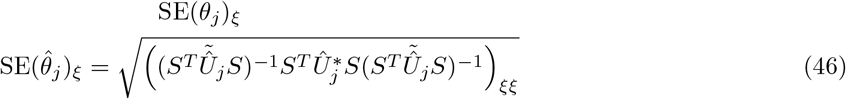

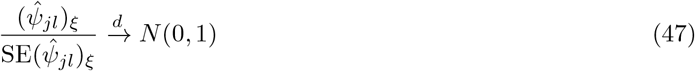

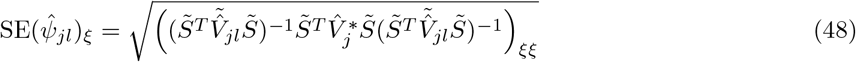

where *ξ* represents the index of the parameter under test in 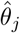 or in 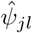. Under asymptotic normality, p-values can easily be calculated using the cumulative distribution function of the standard normal distribution.

#### Non-parametric statistical testing

We observe that our theoretical p-values based on asymptotics are well-calibrated under the null (Supp. Fig. S2). However, this may not always be the case. We therefore consider a second, non-parametric approach for statistical testing. Specifically, for a desired false discovery rate (FDR) *α* ∈ [0, 1], let *t*_1_(*y*), …, *t*_*z*_(*y*) be a set of statistics derived based on the statistics in Eq. (45) and (47), and let *t*_1_(*π*(*y*)), …, *t*_*z*_(*π*(*y*)) be a similar set that was calculated on a permuted *y*, we call *t*_*j*_(*y*) as statistically significant if it is greater than a critical value 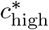, which is the maximal *c* that satisfies:

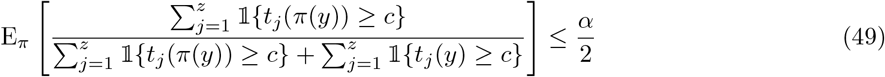

were the expectation is empirically evaluated based on a relatively small number of permutations (but at least ⌈1*/α*⌉). We similarly find the critical value for negative statistics 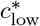, thus obtaining a two-sided test that controls for FDR at level *α*. The critical values can be found by a linear search on the sorted statistics. This approach was previously suggested in the context of a different statistic [30], which had been suggested to suffer from limited power due to possible differences in the null distributions of different genes [31, 32].

We alleviate this risk by standardizing genes by their root mean squares.

### 2.5 Implementation

An R implementation of Keris is available at https://github.com/YosefLab/Keris

## 3 Results

### 3.1 Learning population-level moments from bulk: a simulation study

We first evaluated the capacity of Keris to learn cell-type level moments (means, variances, and gene-gene covariances) from population-level bulk data by mixing gene expression profiles and then evaluating the accuracy of Keris in learning the true moments of the different profiles from the mixtures. To this end, we pooled together expression profiles of different tissues from GTEx [33, 34] and mixed them according to weights that were drawn from an empirical distribution of cell-type proportions that we obtained by applying a reference-based decomposition [35] on a large PBMC data [36]. This resulted in RNA profiles composed of a mixture of several tissues (between 3 and 8), simulating the case of mixtures of cell types.

None of the existing methods in the space of decomposition/deconvolution of genomic data provides estimates for all three moments we considered. In order to establish performance baseline, we therefore considered CIBERSORTx [9] and TCA [24], two deconvolution methods for the estimation of individual-specific cell-type levels (i.e. 3D tensor of samples by genes by cell types) from bulk data. While CIBERSORTx and TCA do not explicitly model or estimate cell-type level gene-gene covariances, we estimated those using the tensors provided by these methods.

We found that CIBERSORTx performs substantially worse than Keris and TCA in estimating all of the three moments even in the simple case of bulk with three cell types (Supp. Fig. S3, S4, and S5); we therefore excluded it from our full benchmarking. TCA performed comparably to Keris in learning the means of the mixed tissues, however, Keris yielded superior performance in estimating variances and covariances (Fig. 3). Importantly, these results were consistent under different sample sizes (n=100 up to n=1000) and using different numbers of tissues for composing the bulk mixtures (Supp Fig. S6, S7)

**Figure 3:**
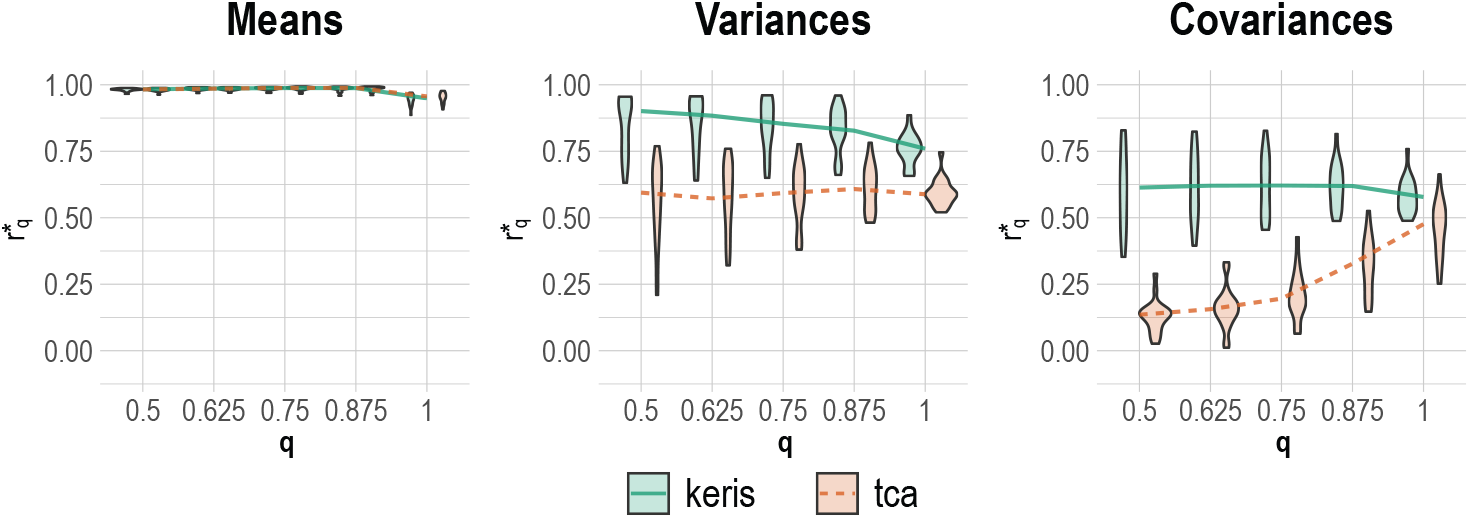
Learning cell-type-specific means, variances, and gene-gene covariances from simulated bulk expression using Keris and TCA. Each of the two methods was applied to mixtures generated from the top *k* = 4 most abundant GTEx tissues: muscle-skeletal (n=801), whole-blood (n=755), skin (n=700), and adipose (n=662). Presented are the results of learning the moments of the tissues composing the mixtures (*n* = 500, *m* = 1000). Results are evaluated using robust correlation 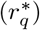 between the true moments of the tissues (estimated directly from the tissues) and the estimated moments, as a function of the data points that were included in the evaluation (*q*; outlier points were excluded based on the joint density of the estimated and true levels). Violin plots represent the performance across 20 different simulations, and the result of a given simulation is the average performance across all four cell types.

### 3.2 Identifying changes in cell-subtype composition with age

We next applied Keris to large bulk PBMC data by Kirsten et al. (n=745) [36] with the goal of identifying genes that are DE with age (below 50 y/o vs. above 70 y/o) in the five immune cell types that contain most of the cells in such samples (monocytes, B, NK, CD8 T, and CD4 T cells). Keris yielded a total of 110 statistically significant DE genes across all cell types (FDR level 0.05). These age-associated changes in a given cell type can result either from change in the composition of cellular sub-types (e.g., naive vs. effector T cells) or from gene expression changes in one or more subsets. To explore this, we inspected the DE genes called by Keris using independent PBMC SC data of four healthy donors (*<*50 y/o or *>*70 y/o) from the Arunachalam et al. study [37]. Indeed, using the Arunachalam data, we found that 48 out of the 110 associations returned by Keris are marker genes for subtypes of the five cell types we modeled. Concretely, the gene *SNX22*, which Keris estimated as upregulated with age in B cells in the Kirsten bulk data, turned out to be upregulated in intermediate B cells compared with other B cell subtypes in the Arunachalam data (p-value=3.1e-05; Wilcoxon test). This therefore suggests an increase in composition of intermediate B cells with age. Importantly, we were able to further confirm this result using data from 70 individuals across two independent PBMC SC studies by Stephenson et al. [15] and Ren et al. [16] (p-value=0.034; linear regression of intermediate B cell composition on age, while accounting for disease condition, sex, and batch). Similarly, the gene *LRRN3*, which Keris called as downregulated with age in CD4 T cells, turned out to be upregulated in naive CD 4 T cells compared with other subtypes of CD4 T cells (p-value=2.7e-20; Wilcoxon test). This suggests a decrease in CD4 naive T cells with age, a well-known result [38] that we were able to further validate in the Stephenson and Ren datasets (p-value=0.031, linear regression of CD4 naive T cell composition on age, while accounting for disease condition, sex, and batch).

In the monocytes class, Keris reported *LGALS3* as downregulated with age, which, in conjunction with the gene’s downregulation in CD16 monocytes compared with CD14 monocytes in the Arunachalam data (p-value=7.2e-19; Wilcoxon test), suggests an increase in CD16 monocyte composition with age. We could not detect evidence for this result in the Stephenson and Ren datasets (p-value=0.33; linear regression), however, we note that an age-associated increase in CD16 monocyte composition was previously reported in a large study with PBMC [39]. Lastly, the rest of the genes that were reported by Keris as DE and found to be marker genes for cell subtypes in the SC data are strong markers for more than one cell subtype. Hence, we were not able to conclusively attribute them to changes in composition of particular cell subtypes.

### 3.3 Capturing senescent cells via Keris-integrated senescence score

We expected the 62 Keris-derived DE genes that were not identified as cell-subtype markers in the Arunachalam data to reflect changes in expression rather than changes in composition with age. Following our model, we expected these DE genes to capture effects in populations of aging (senescent) cells. Clearly, single genes may demonstrate small effects with age. Therefore, given multiple DE genes in a particular cell type, we created a Keris-integrated senescence score (KISS) for integrating over the effects coming from all the Keris-derived DE genes in that cell type. Specifically, we defined the KISS value of a single cell to be the total expression across all age-upregulated DE genes minus the expression across all age-downregulated DE genes. Since most of the 62 Keris-derived genes that were expected to reflect changes in expression were called as DE in either NK or CD8 T cells (39 and 19 genes, respectively), we focused only on these cell types.

We first calculated an NK-specific KISS for every NK cell in healthy PBMC samples (*<*50 y/o or *>*70 y/o) from the Stephenson (n=23) [15] and Ren (n=20) [16] datasets. In order to confirm that high KISS levels indeed tag senescent cells, we considered *CDKN2A*, a known marker gene of cellular senescence [40]. While *CDKN2A* is not considered as a very accurate marker for cellular senescence, we hypothesized that cells expressing *CDKN2A* (denote *CDKN2A*+, as opposed to *CDKN2A*-) will be enriched with higher KISS levels. Indeed, we found that *CDKN2A*+ NK cells demonstrate heavier tail of high NK-specific KISS values compared with *CDKN2A*-NK cells in both the Stephenson (p-value=1.9e-4) and Ren (p-value=6e-4) datasets; p-values were calculated by comparing the top quartile of KISS in *CDKN2A*+ cells with 10^5^ equally-sized random subsets of *CDKN2A*-cells. While we do not expect all cells coming from aged individuals to necessarily be senescent, a reasonable cellular senescence score is expected to distribute somewhat differently in cells from aged individuals compared with cells from young individuals. As expected, cells from aged individuals (*>*70 y/o) demonstrated higher NK-specific KISS values compared with cells from younger individuals (*<*50 y/o; Stephenson p-value=5.2e-3, Ren p-value=1.5e-12; t-test).

Repeating the above analysis for CD8 T cells, we found that *CDKN2A*+ CD8 T cells are not enriched for high CD8-specific KISS values in both the Stephenson and Ren datasets (p-value=1). However, we suggest that *CDKN2A* may not be a very informative marker of senescence in CD8 T cells for two reasons. First, CD8 T cells from aged individuals do demonstrate higher CD8-specific KISS values compared with cells from younger individuals (Stephenson p-value=0.017, Ren p-value *<*2.2e-16; t-test; and Fig. 4). Second, we observe that aged individuals have more extreme CD8-specific KISS values in their CD8 T cells when compared to young individuals. Specifically, for every individual sample in the Stephenson and Ren datasets, we considered its top decile of the KISS values (i.e., across all CD8 T cells of the individual), and we found that all young individuals except for one had a top decile value of zero, while the aged individuals in the data (n=2) had a positive top decile. Interestingly, we did not observe a similar trend in the population of NK cells using the NK-specific KISS, suggesting either a limited accuracy of our score for NK cells or the need for larger sample sizes for evaluation. Henceforth, we focused on our CD8-specific KISS.

**Figure 4:**
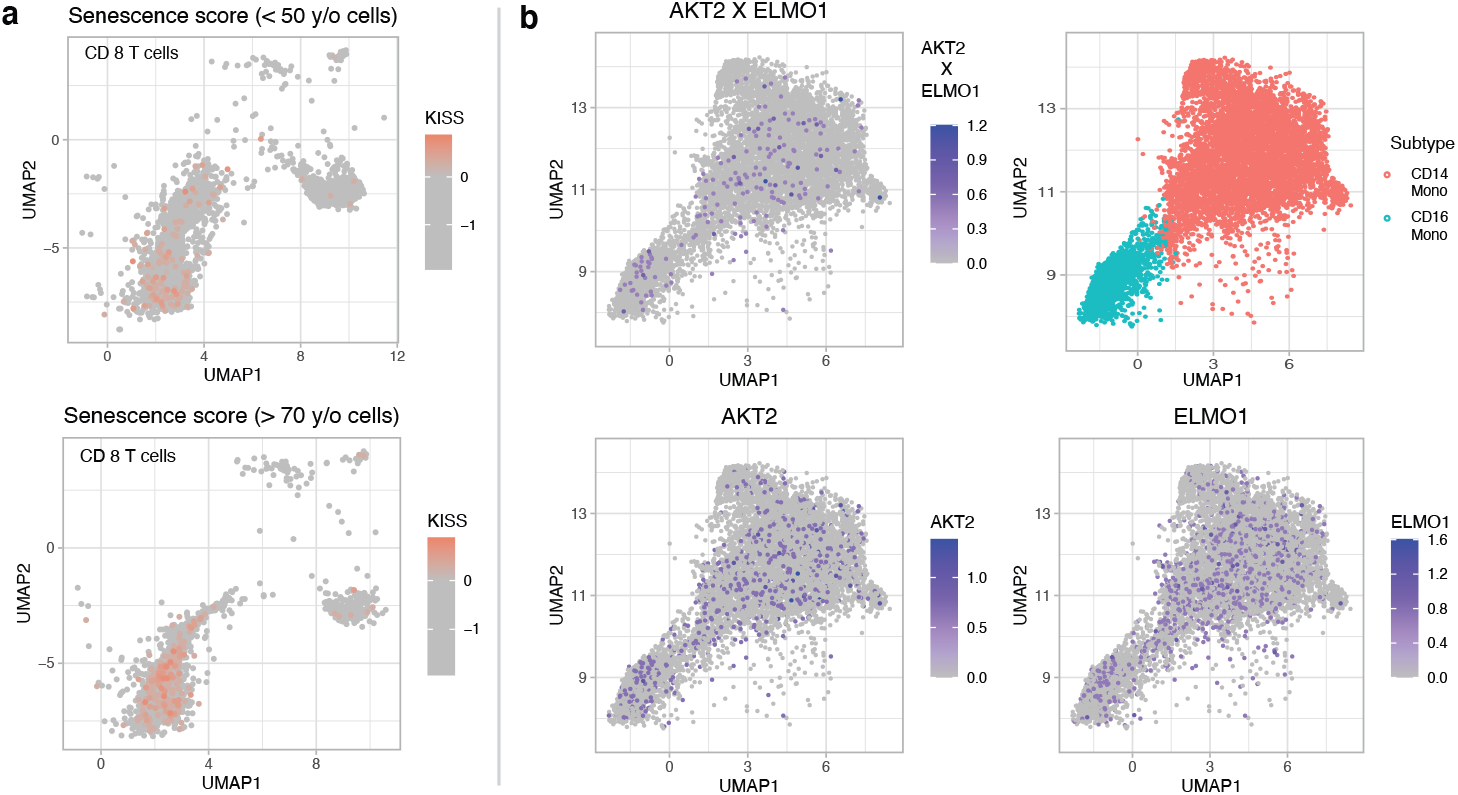
(a) CD8-specific KISS levels in CD8 T cells from young (top) and aged (bottom) individuals in the Ren data; cells were randomly subsampled to match in numbers and plotted using UMAPs [41]. (b) The expression pattern of the interaction *AKT2* X *ELMO1* in monocyte cells is not clearly reflected in the separate expression of the genes *AKT2* and *ELMO1*; presented are UMAPs based on the Stephenson data.

Age-related effects on expression are expected to be heterogeneous, as reflected by the inconsistent calling of genes that are with DE age across small studies (Supp. Fig. S1). Since our definition of KISS is based on DE genes detected in population-level data, we expected that comparing CD8 T cells with high KISS levels to CD8 T cells with low KISS levels will demonstrate differences that are consistent across studies. Indeed, performing a DE analysis (Wilcoxon test) with low-KISS cells (first quartile of KISS values) versus high-KISS cells (fourth quartile) revealed high consistency between the Stephenson and Ren data (J=0.61; Jaccard index). This result is particularly remarkable due to the fact that it relies on a single aged individual (*>*70 y/o) in each of the two datasets. In contrast, taking a standard approach of performing DE analysis based on cells from young individuals versus cells from aged individuals yielded poor consistency (J=0.14). Finally, this gap in consistency was further manifested in a gene enrichment analysis based on the DE results. Considering the top 10 most positively enriched GO term and the top 10 most negatively enriched terms revealed a much higher overlap between the Stephenson and Ren datasets when usign the Keris-derived KISS values (17 GO terms out of 20) compared with the standard approach for DE analysis (10 GO terms).

### 3.4 Applying Keris to detect cell-type level gene-gene interactions

Lastly, we applied Keris again to the Kirsten bulk PBMC data, this time with the goal to demonstrate the ability of Keris to detect cell-type level differential correlations (DC). Specifically, we aimed at detecting age-associated DC in monocyte cells, while considering only the group of genes that are tagged by KEGG [42] as involved in either chemokine signaling or phagocytosis, two cellular pathways that are known to demonstrate altered function with age in monocytes [43] (a total of 417 genes). Keris reported two pairs of genes as significantly DC with age in monocytes (based on permutation test with 10^5^ permutations). Concretely, Keris estimated the correlation between *DOCK2* and *CX3CR1* and the correlation between *AKT2* and *ELMO1* to be upregulated in monocytes. Using the Arunachalam SC data, we found the interaction *DOCK2* X *CX3CR1* to be upregulated in CD16 monocytes compared with CD14 monocytes (p-value*<*2.2e-16; t-test), thus reinforcing the previous indication by Keris of an increase in CD16 monocyte composition with age (based on *LGALS3*). Since we did not observe the interaction *AKT2* X *ELMO1* to be a notable marker for monocyte subtypes (p-value=0.04; t-test), we evaluated whether this interaction captures senescent cells. We could not find evidence that *CDKN2A*+ monocyte cells are enriched for high *AKT2* X *ELMO1* levels (p-value *>*0.14 in both Stephenson and Ren datasaets; based on 10^5^ equally-sized random subsets of *CDKN2A*-cells). However, similarly to our analysis with the DE results, we found evidence that monocyte cells from aged individuals demonstrate *AKT2* X *ELMO1* more frequently than monocyte cells from young individuals (Stephenson p-value=0.001, Ren p-value=0.066; t-test). Importantly, the cells tagged by the interaction *AKT2* X *ELMO1* could not be revealed by looking at either *AKT2* or *ELMO1* separately (Fig. 4), thus showing the promise of considering gene-gene interactions for profiling populations of senescent cells.

## 4 Discussion

We suggest that results returned by Keris for a particular cell type can be attributed to changes in cell-subtype composition if they are markers for known sub-populations of that cell type. However, it is possible that in some cases marker genes for cell subtypes will also demonstrate changes in expression with the condition under test. Since it is not clear how such signals can be disentangled, in this a scenario we are bound to ignore such genes in our downstream analysis of changes in expression. However, given the high consistency between cell-type markers across individuals (e.g., Fig. 1), we expect these cases to be infrequent.

Finally, our work focuses on SC and bulk expression data, however, all of our assumptions are distribution-free. Hence, in theory, Keris is expected to be applicable to other genomic modalities as well.

## Supplementary Figures

**Figure S1:**
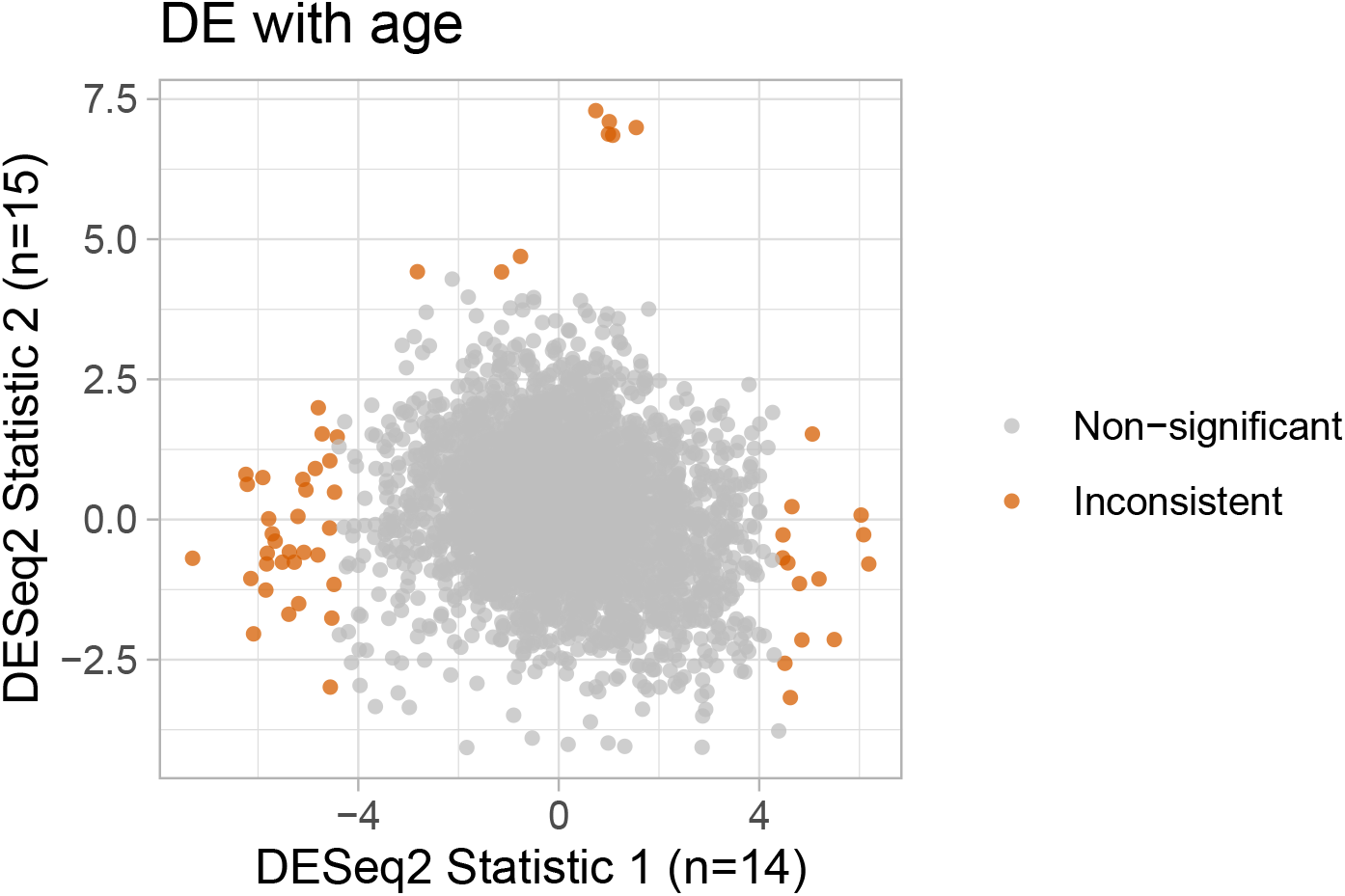
Evaluating the consistency of differentially expressed (DE) genes with age across two independent PBMC SC datasets [1, 2]. Presented are the test statistics of DE analysis between young (n=14; *<*50 y/o) and aged (n=14; *>*70 y/o) healthy individuals, *within* each of the main immune cell types (results from all cell types were pooled together). Consistency was defined as the Jaccard index – the fraction of consistently DE genes (significant in both datasets) out of the total number of DE genes (significant at least once) using DESeq2 [3] on the top 1000 most highly expressed genes, while accounting for sex.

**Figure S2:**
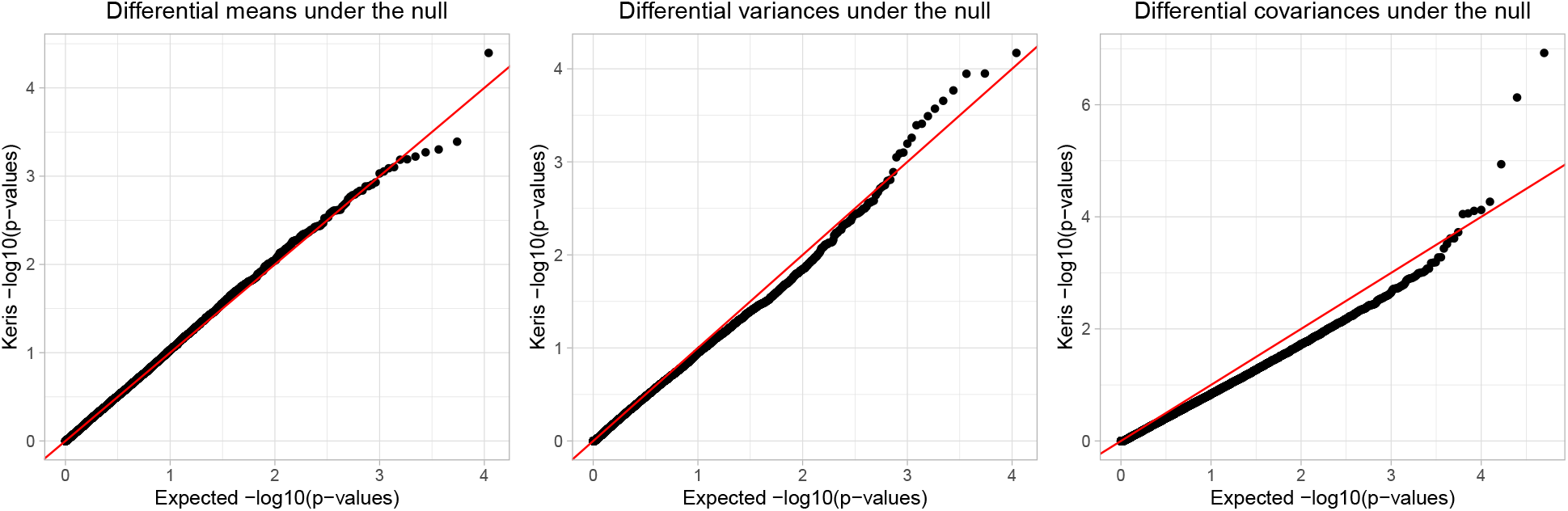
Distribution of p-values under the asymptotic distribution of the Keris statistic in the Kirsten et al. data [4] after permuting the condition label (*<*50 y/o or *>*70 y/o). Presented at the distribution of p-values under the null for differential means, variances, and correlations (10,000 randomly selected pairs of genes); distributions were pooled across cell types.

**Figure S3:**
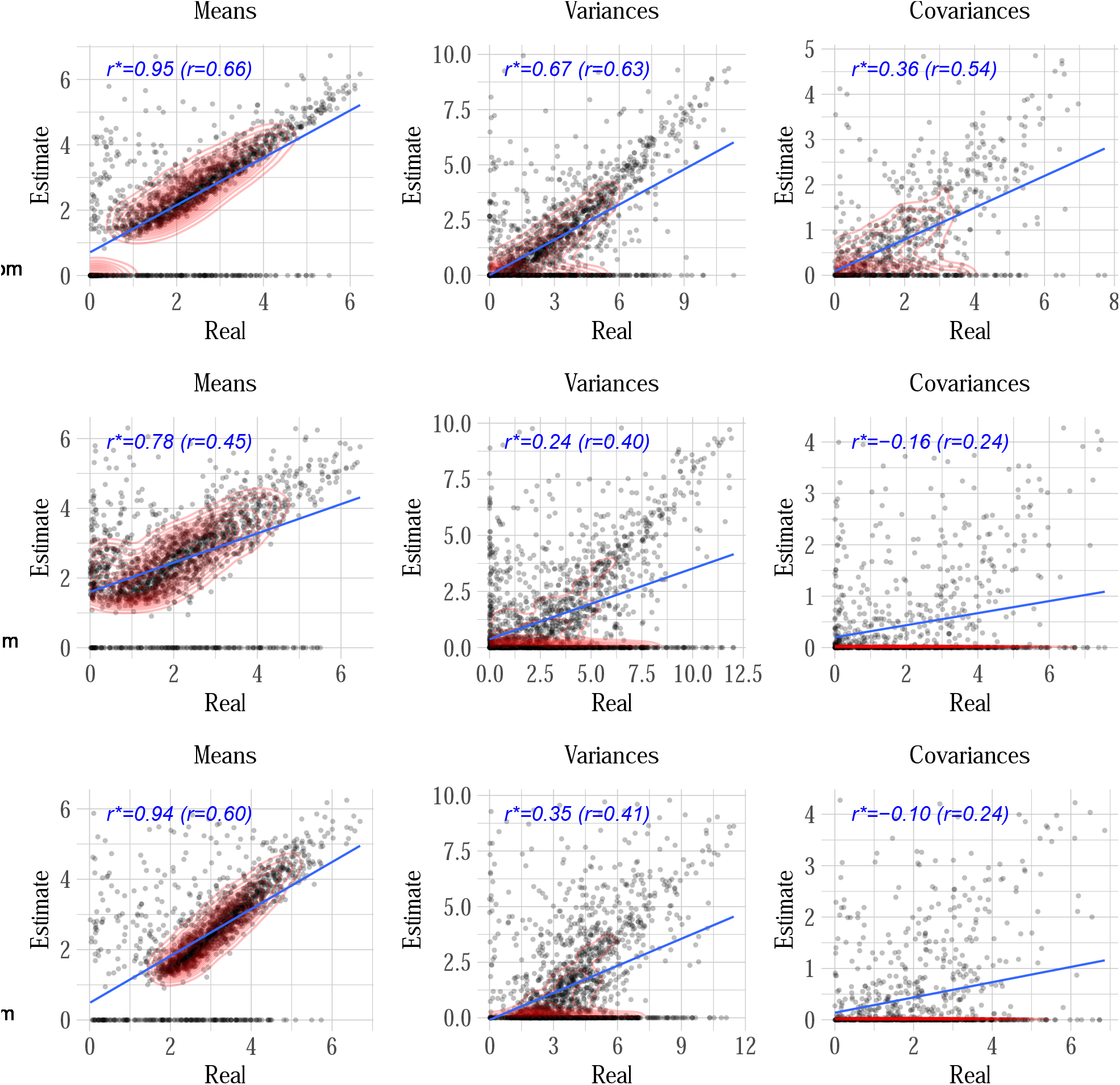
Learning cell-type-specific means, variances, and gene-gene covariances from simulated bulk expression using CIBERSORTx. Mixtures were generated from the top *k* = 3 most abundant GTEx tissues: muscle-skeletal (n=801), whole-blood (n=755), and skin (n=700). Presented are the results of learning the moments of the tissues composing the mixtures (*n* = 500, *m* = 1000). Results were evaluated for each tissue (in a separate row) using robust correlation (*r*^*^) between the true moments of the tissues (estimated directly from the tissues) and the CIBERSORTx estimates of the moments. Two-dimensional density plots colored in red show the majority of points (75%), based on which a robust correlation was calculated in each plot.

**Figure S4:**
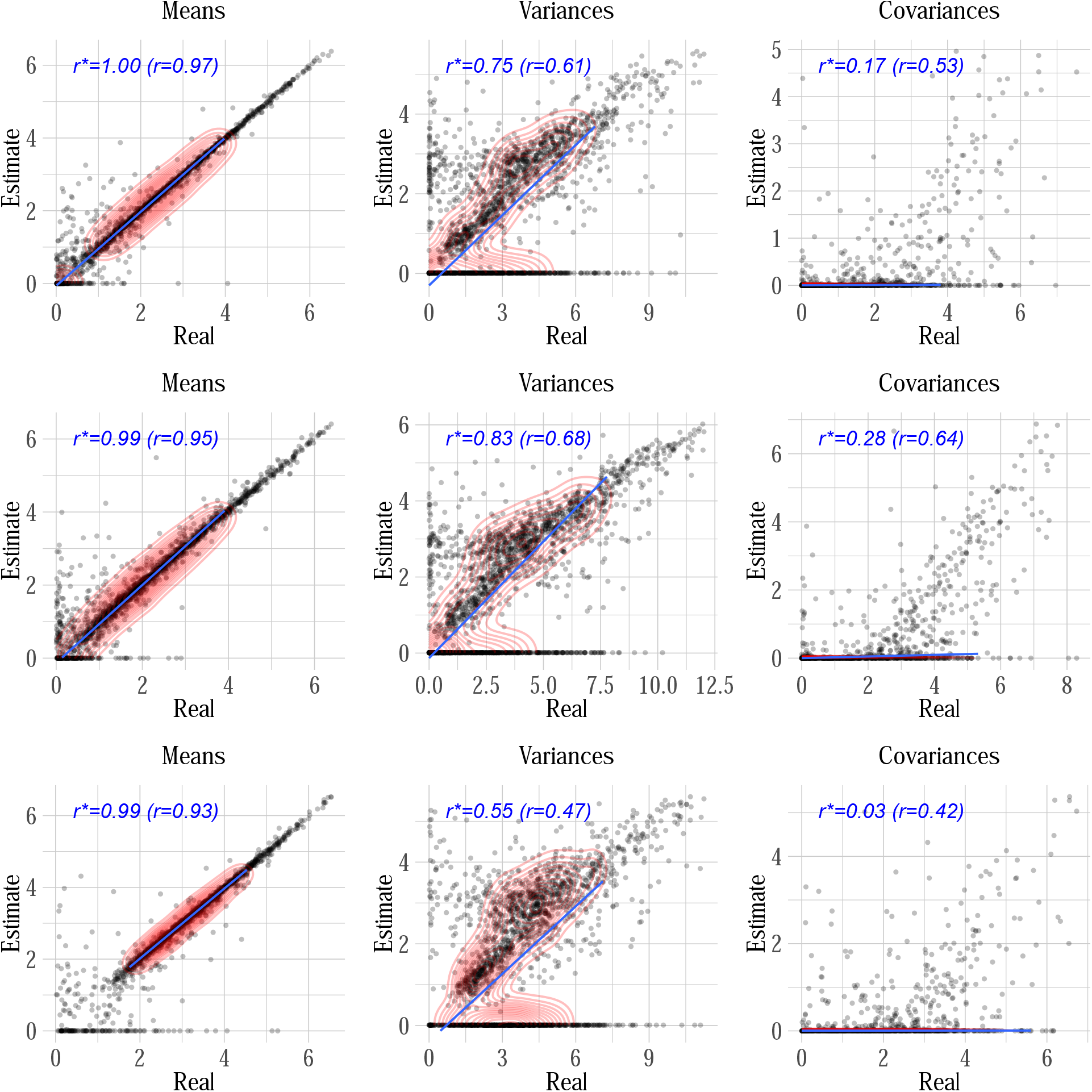
Learning cell-type-specific means, variances, and gene-gene covariances from simulated bulk expression using TCA. Mixtures were generated from the top *k* = 3 most abundant GTEx tissues: muscle-skeletal (n=801), whole-blood (n=755), and skin (n=700). Presented are the results of learning the moments of the tissues composing the mixtures (*n* = 500, *m* = 1000). Results were evaluated for each tissue (in a separate row) using robust correlation (*r*^*^) between the true moments of the tissues (estimated directly from the tissues) and the TCA estimates of the moments. Two-dimensional density plots colored in red show the majority of points (75%), based on which a robust correlation was calculated in each plot.

**Figure S5:**
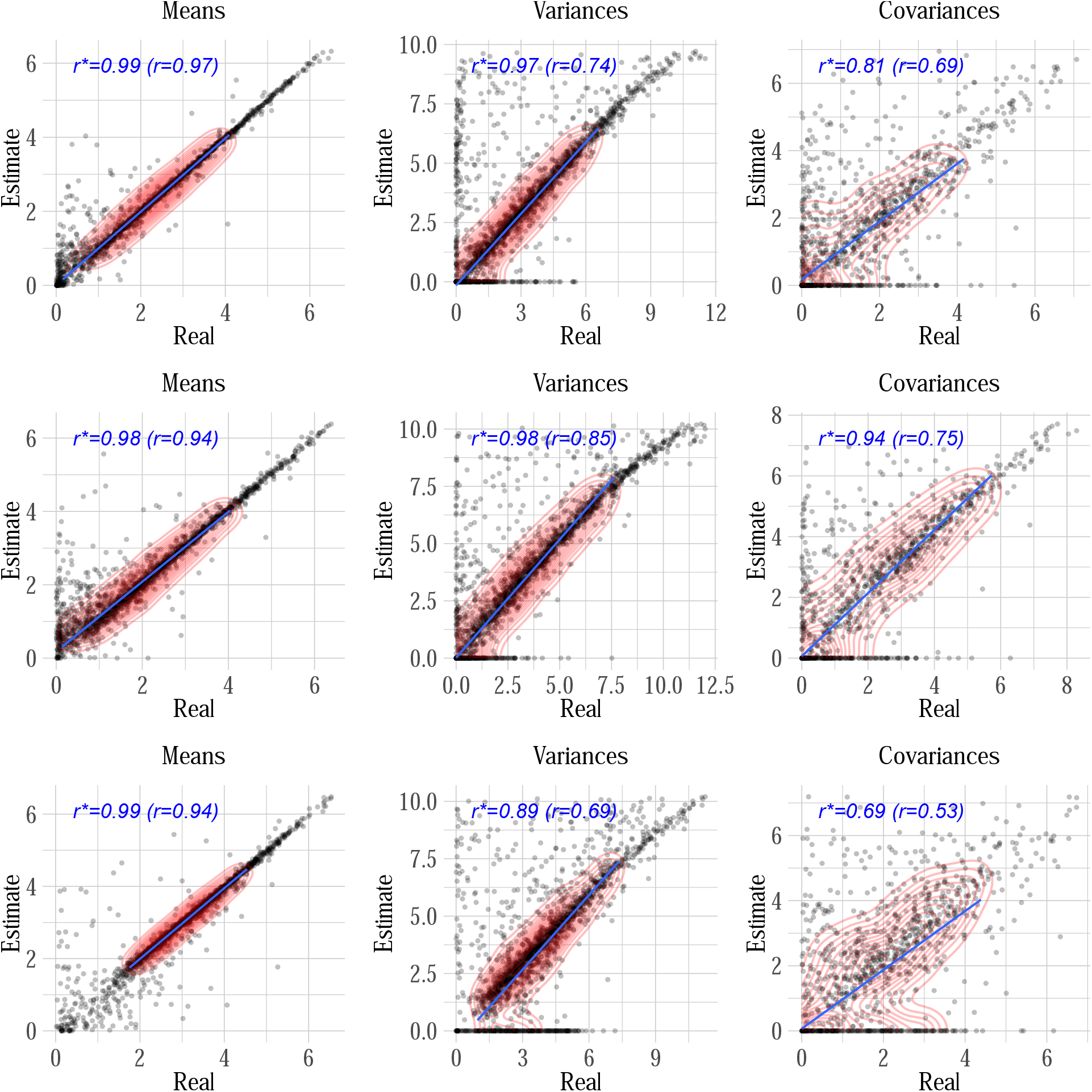
Learning cell-type-specific means, variances, and gene-gene covariances from simulated bulk expression using Keris. Mixtures were generated from the top *k* = 3 most abundant GTEx tissues: muscle-skeletal (n=801), whole-blood (n=755), and skin (n=700). Presented are the results of learning the moments of the tissues composing the mixtures (*n* = 500, *m* = 1000). Results were evaluated for each tissue (in a separate row) using robust correlation (*r*^*^) between the true moments of the tissues (estimated directly from the tissues) and the Keris estimates of the moments. Two-dimensional density plots colored in red show the majority of points (75%), based on which a robust correlation was calculated in each plot.

**Figure S6:**
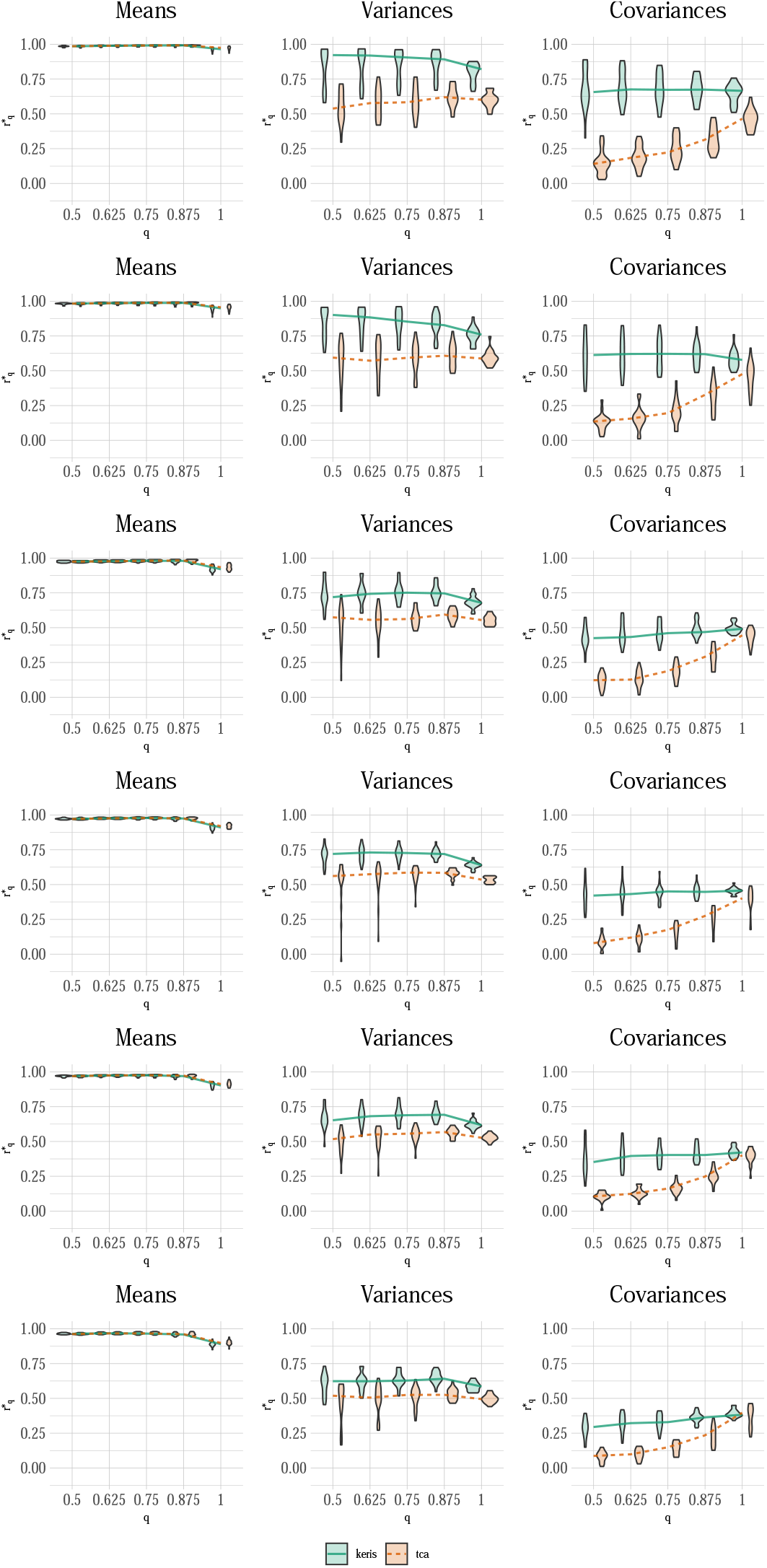
Learning cell-type-specific means, variances, and gene-gene covariances from simulated bulk expression using Keris and TCA. Each of the two methods was applied to mixtures generated from 3 to 8 most abundant GTEx tissues (first row corresponds to *k* = 3 tissues, the second row corresponds to *k* = 4 tissues and so on). Presented are the results of learning the moments of the tissues composing the mixtures (*n* = 500, *m* = 1000). Results are evaluated using robust correlation 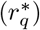 between the true moments of the tissues (estimated directly from the tissues) and the estimated moments, as a function of the data points that were included in the evaluation (*q*; outlier points were excluded based on the joint density of the estimated and true levels). Violin plots represent the performance across 20 different simulations, and the result of a given simulation is the average performance across all cell types.

**Figure S7:**
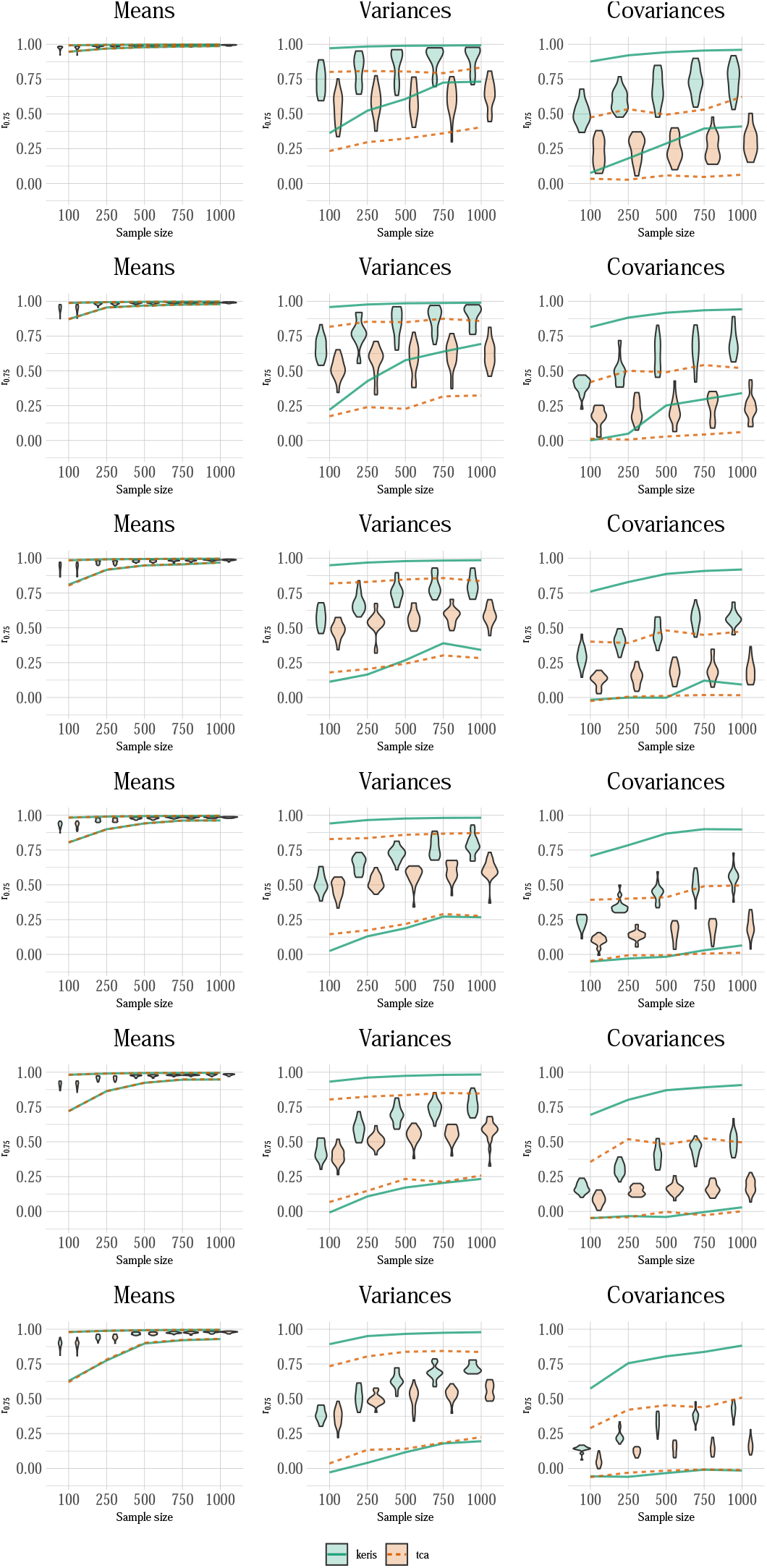
Learning cell-type-specific means, variances, and gene-gene covariances from different dataset sizes of simulated bulk expression using Keris and TCA. Each of the two methods was applied to mixtures generated from 3 to 8 most abundant GTEx tissues (first row corresponds to *k* = 3 tissues, the second row corresponds to *k* = 4 tissues and so on). Presented are the results of learning the moments of the tissues composing the mixtures (*n* = 500, *m* = 1000). Results are evaluated using robust correlation (*r*^*^) between the true moments of the tissues (estimated directly from the tissues) and the estimated moments, as a function of the number of simulated samples. Violin plots represent the performance across 20 different simulations, the result of a given simulation is the average performance across all cell types, as solid and dashed lines represent the median of the best and worse performing cell type across all 20 simulations.

## References

[1] Roser Vento-Tormo et al. “Single-cell reconstruction of the early maternal–fetal interface in humans”. In: Nature 563.7731 (2018), pp. 347–353.

[2] Zhilei Bian et al. “Deciphering human macrophage development at single-cell resolution”. In: Nature 582.7813 (2020), pp. 571–576.

[3] Chiara Baccin et al. “Combined single-cell and spatial transcriptomics reveal the molecular, cellular and spatial bone marrow niche organization”. In: Nature cell biology 22.1 (2020), pp. 38–48.

[4] Massimo Mangino et al. “Innate and adaptive immune traits are differentially affected by genetic and environmental factors”. In: Nature communications 8.1 (2017), pp. 1–7.

[5] Ron Edgar, Michael Domrachev, and Alex E Lash. “Gene Expression Omnibus: NCBI gene expression and hybridization array data repository”. In: Nucleic acids research 30.1 (2002), pp. 207–210.

[6] Maayan Baron et al. “A single-cell transcriptomic map of the human and mouse pancreas reveals inter-and intra-cell population structure”. In: Cell systems 3.4 (2016), pp. 346–360.

[7] Lingxue Zhu et al. “A unified statistical framework for single cell and bulk RNA sequencing data”. In: The annals of applied statistics 12.1 (2018), p. 609.

[8] Xuran Wang et al. “Bulk tissue cell type deconvolution with multi-subject single-cell expression reference”. In: Nature communications 10.1 (2019), pp. 1–9.

[9] Aaron M Newman et al. “Determining cell type abundance and expression from bulk tissues with digital cytometry”. In: Nature biotechnology 37.7 (2019), pp. 773–782.

[10] Brandon Jew et al. “Accurate estimation of cell composition in bulk expression through robust integration of single-cell information”. In: Nature communications 11.1 (2020), pp. 1–11.

[11] Meichen Dong et al. “SCDC: bulk gene expression deconvolution by multiple single-cell RNA sequencing references”. In: Briefings in bioinformatics 22.1 (2021), pp. 416–427.

[12] Pawel F Przytycki and Katherine S Pollard. “CellWalker integrates single-cell and bulk data to resolve regulatory elements across cell types in complex tissues”. In: Genome biology 22.1 (2021), pp. 1–16.

[13] Wolf Herman Fridman et al. “The immune contexture in human tumours: impact on clinical outcome”. In: Nature Reviews Cancer 12.4 (2012), pp. 298–306.

[14] Jacques Rahier, RM Goebbels, and Jean-Claude Henquin. “Cellular composition of the human diabetic pancreas”. In: Diabetologia 24.5 (1983), pp. 366–371.

[15] Emily Stephenson et al. “Single-cell multi-omics analysis of the immune response in COVID-19”. In: Nature medicine 27.5 (2021), pp. 904–916.

[16] Xianwen Ren et al. “COVID-19 immune features revealed by a large-scale single-cell transcriptome atlas”. In: Cell 184.7 (2021), pp. 1895–1913.

[17] Michael I Love, Wolfgang Huber, and Simon Anders. “Moderated estimation of fold change and dispersion for RNA-seq data with DESeq2”. In: Genome biology 15.12 (2014), pp. 1–21.

[18] Peter Langfelder and Steve Horvath. “WGCNA: an R package for weighted correlation network analysis”. In: BMC bioinformatics 9.1 (2008), pp. 1–13.

[19] Shila Ghazanfar et al. “DCARS: differential correlation across ranked samples”. In: Bioinformatics 35.5 (2019), pp. 823–829.

[20] Marjan Farahbod and Paul Pavlidis. “Untangling the effects of cellular composition on coexpression analysis”. In: Genome research 30.6 (2020), pp. 849–859.

[21] Shahin Mohammadi et al. “A critical survey of deconvolution methods for separating cell types in complex tissues”. In: Proceedings of the IEEE 105.2 (2016), pp. 340–366.

[22] Jiebiao Wang, Bernie Devlin, and Kathryn Roeder. “Using multiple measurements of tissue to estimate subject-and cell-type-specific gene expression”. In: Bioinformatics 36.3 (2020), pp. 782–788.

[23] Yun Zhang et al. “The effect of tissue composition on gene co-expression”. In: Briefings in bioinformatics 22.1 (2021), pp. 127–139.

[24] Elior Rahmani et al. “Cell-type-specific resolution epigenetics without the need for cell sorting or single-cell biology”. In: Nature communications 10.1 (2019), pp. 1–11.

[25] Ana Conesa et al. “A survey of best practices for RNA-seq data analysis”. In: Genome biology 17.1 (2016), pp. 1–19.

[26] Davide Risso et al. “Normalization of RNA-seq data using factor analysis of control genes or samples”. In: Nature biotechnology 32.9 (2014), pp. 896–902.

[27] Oliver Stegle et al. “Using probabilistic estimation of expression residuals (PEER) to obtain increased power and interpretability of gene expression analyses”. In: Nature protocols 7.3 (2012), pp. 500–507.

[28] Lars Peter Hansen. “Large sample properties of generalized method of moments estimators”. In: Econometrica: Journal of the econometric society (1982), pp. 1029–1054.

[29] Fumio Hayashi. “Econometrics,? Princeton University Press: Princeton”. In: (2000).

[30] Virginia Goss Tusher, Robert Tibshirani, and Gilbert Chu. “Significance analysis of microarrays applied to the ionizing radiation response”. In: Proceedings of the National Academy of Sciences 98.9 (2001), pp. 5116–5121.

[31] Wei Pan. “On the use of permutation in and the performance of a class of nonparametric methods to detect differential gene expression”. In: Bioinformatics 19.11 (2003), pp. 1333–1340.

[32] Yang Xie, Wei Pan, and Arkady B Khodursky. “A note on using permutation-based false discovery rate estimates to compare different analysis methods for microarray data”. In: Bioinformatics 21.23 (2005), pp. 4280–4288.

[33] Latarsha J Carithers et al. “A novel approach to high-quality postmortem tissue procurement: the GTEx project”. In: Biopreservation and biobanking 13.5 (2015), pp. 311–319.

[34] GTEx Consortium et al. “The Genotype-Tissue Expression (GTEx) pilot analysis: Multitissue gene regulation in humans”. In: Science 348.6235 (2015), pp. 648–660.

[35] Francesca Finotello et al. “Molecular and pharmacological modulators of the tumor immune contexture revealed by deconvolution of RNA-seq data”. In: Genome medicine 11.1 (2019), pp. 1–20.

[36] Holger Kirsten et al. “Dissecting the genetics of the human transcriptome identifies novel trait-related trans-eQTLs and corroborates the regulatory relevance of non-protein coding loci”. In: Human molecular genetics 24.16 (2015), pp. 4746–4763.

[37] Prabhu S Arunachalam et al. “Systems biological assessment of immunity to mild versus severe COVID-19 infection in humans”. In: Science 369.6508 (2020), pp. 1210–1220.

[38] Nels C Olson et al. “Decreased naive and increased memory CD4+ T cells are associated with subclinical atherosclerosis: the multi-ethnic study of atherosclerosis”. In: PloS one 8.8 (2013), e71498.

[39] Sebastian Seidler et al. “Age-dependent alterations of monocyte subsets and monocyte-related chemokine pathways in healthy adults”. In: BMC immunology 11.1 (2010), pp. 1–11.

[40] Janakiraman Krishnamurthy et al. “Ink4a/Arf expression is a biomarker of aging”. In: The Journal of clinical investigation 114.9 (2004), pp. 1299–1307.

[41] Leland McInnes, John Healy, and James Melville. “Umap: Uniform manifold approximation and projection for dimension reduction”. In: arXiv preprint 1802.03426 (2018).

[42] Minoru Kanehisa and Susumu Goto. “KEGG: kyoto encyclopedia of genes and genomes”. In: Nucleic acids research 28.1 (2000), pp. 27–30.

[43] Tamas Fulop et al. Handbook of Immunosenescence. Springer, 2019.

## References

[1] Emily Stephenson et al. “Single-cell multi-omics analysis of the immune response in COVID-19”. In: Nature medicine 27.5 (2021), pp. 904–916.

[2] Xianwen Ren et al. “COVID-19 immune features revealed by a large-scale single-cell transcriptome atlas”. In: Cell 184.7 (2021), pp. 1895–1913.

[3] Michael I Love, Wolfgang Huber, and Simon Anders. “Moderated estimation of fold change and dispersion for RNA-seq data with DESeq2”. In: Genome biology 15.12 (2014), pp. 1–21.

[4] Holger Kirsten et al. “Dissecting the genetics of the human transcriptome identifies novel trait-related trans-eQTLs and corroborates the regulatory relevance of non-protein coding loci”. In: Human molecular genetics 24.16 (2015), pp. 4746–4763.

